# A Haploid Genetic Screening Method for Proteins Influencing Mammalian Nonsense-Mediated mRNA Decay Activity

**DOI:** 10.1101/452490

**Authors:** Maximilian W. Popp, Lynne E. Maquat

## Abstract

Despite a long appreciation for the role of nonsense-mediated mRNA decay (NMD) in the destruction of faulty, disease-causing mRNAs, as well as its role in the maintenance of normal, endogenous transcript abundance, systematic unbiased methods for uncovering modifiers of NMD activity in mammalian cells remain scant. Here we present and validate a haploid genetic screening method for identifying proteins and processes that stimulate NMD activity involving a 3′-untranslated region exon-junction complex. This reporter-based screening method can be adapted for interrogating other pathways whose output can be measured by the intracellular production of fluorescent proteins.

## Introduction

Nonsense-mediated mRNA decay (NMD) is a cellular quality control mechanism used to inspect and destroy aberrant mRNAs that have the potential to code for toxic truncated proteins (1). The best characterized mechanism for triggering NMD involves transcripts that derive from genes containing a premature termination codon (PTC) upstream of an exon-exon junction, a configuration that is aberrant, since most mammalian genes contain the termination codon in their last exon. During the process of pre-mRNA splicing, a large macromolecular complex called the exon junction complex (EJC) is deposited slightly (~20-25-nucleotides) upstream from the spliced exon-exon junction. When this junction occurs >50-55 nucleotides downstream of the PTC, in what is effectively the 3′-untranslated region (3′UTR), the size of the ribosome is insufficient to remove the EJC during translation, a signal that identifies the mRNA for NMD. Briefly, a set of core factors involving the proteins UP-Frameshift (UPF) 1 and Suppressor with Morphological effect on Genitalia 1 (SMG1) assembles at the terminating ribosome. Generally, the EJC anchors the protein UPF3X (also called UPF3B), which interacts with and displays UPF2 to the complex at the terminating ribosome. When these two protein complexes interact (a situation that cannot occur without a 3′UTR EJC), an ill-defined step occurs that culminates in SMG1-mediated phosphorylation of UPF1. This step is the commitment step to NMD and, if triggered, results in destruction of the mRNA. Phosphorylated UPF1 recruits the endonuclease SMG6 as well as the SMG5/7 complex, which subsequently recruits decapping and deadenylating activities to degrade the aberrant transcript (1). Other normal, non-mutated transcript features such as an unusually long 3′UTR, a 5′UTR intron, the presence of a UGA selenocysteine-encoding codon, or alternative splicing (AS) events that purposefully locate a PTC >50-55 nucleotides upstream of an exon-exon junction allow the cell to exploit the NMD system to control an estimated ~10% of normal cellular transcripts. Cells seem to do this to allow for adaptation to changing environments (2-9).

Additional proteins have more recently been discovered that can modulate the activity of mammalian NMD. For example, the SMG8/9 proteins (10) inhibit the kinase activity of SMG1. DExH-Box Helicase 34 (DHX34) and Neuroblastoma Amplified Sequence (NBAS) co-regulate NMD sensitive transcripts, cooperating with the core NMD factors (11,12). The splicing protein CWC22 couples splicing to EJC deposition, facilitating NMD (13). Interactor of little elongation Complex ELL subunit 1 (ICE1) facilitates anchoring of UPF3X to the EJC (14), and Polypyrimidine Tract Binding Protein 1 (PTBP1) shields some transcripts with long 3′UTRs from NMD (15). Cellular pathways can also influence NMD activity by incompletely understood mechanisms. For example, a rise in intracellular calcium levels has been shown to inhibit NMD (16). Initiation of the unfolded protein response (5,17) and induction of the general stress response also inhibit NMD (2,3,5,9). Certain developmental transitions require adjustment of NMD activity (18,19), and induction of apoptosis is associated with the attenuation of NMD (4,7,8).

Genetic methods for identifying new pathways and proteins that modulate NMD activity remain few. A recent CRISPR-based screen combined with a highly engineered reporter system identified several potential new NMD regulators (20), and a short interfering RNA (siRNA)-based screen identified ICE1 as an NMD regulator (14). Both screening methods are confined to preplanned targeting of genes to which the appropriate reagents (guide RNAs or siRNAs) can be designed *in silico*. siRNAs are also known to incompletely disrupt target gene function. Here, we present an adaptation of a well-documented method that uses unbiased targeting of a mutagen (a gene-trap retrovirus) applied to a haploid mammalian cell line (21-30) engineered to accurately report on NMD activity in order to screen for genes and pathways that enhance NMD activity.

## Results

### Reporter Design Considerations and Delivery

We considered several important criteria when designing a reporter that accurately reflects the activity of NMD (**Fig. 1A**). First, the reporter should have the necessary cis-information that renders it susceptible to NMD. In the case of 3′UTR EJC-enhanced NMD, this takes the form of an intron that violates the 50-55-nucleotide rule—the intron should be located >50-55-nucleotides downstream of a termination codon to be interpreted as aberrant. Because the T-cell receptor ß (TCRß) minigene mRNA is a well-characterized NMD substrate (31), we used a portion of this transcript including the JC intron (the intron between the joining J segment and the constant C segment). Second, we considered the read-out of our screen. Aiming to use flow cytometry, i.e. fluorescence activated cell sorting (FACS) to identify changes in NMD, we placed the coding region of an N-terminally FLAG-tagged mCherry protein upstream of the TCRß minigene fragment. The natural stop codon should thus mimic a premature termination codon (PTC) since it resides >50-nucleotides upstream of the JC intron. The use of the natural stop codon is important since the small amount of protein that is produced from the NMD substrate should contain no traces of TCRß-derived protein. This is because HAP1, which were derived from the male chronic myelogenous leukemia KBM-7 cell line (24), do not contain TCRα or TCRß proteins, and unpaired TCR chains are well-characterized subjects of dislocation from the endoplasmic reticulum followed by rapid proteosomal degradation (32), a condition that would limit signal in our FACS-based assay. Finally, as a control and to facilitate downstream steps (see below), we flanked the TCRß sequences with locus of X-over P1 (loxP) sites recognized by the Cre recombinase. As an additional control, we generated a version of our reporter that lacks only the JC intron (ΔJC) and is therefore immune to NMD.

**Figure 1.**
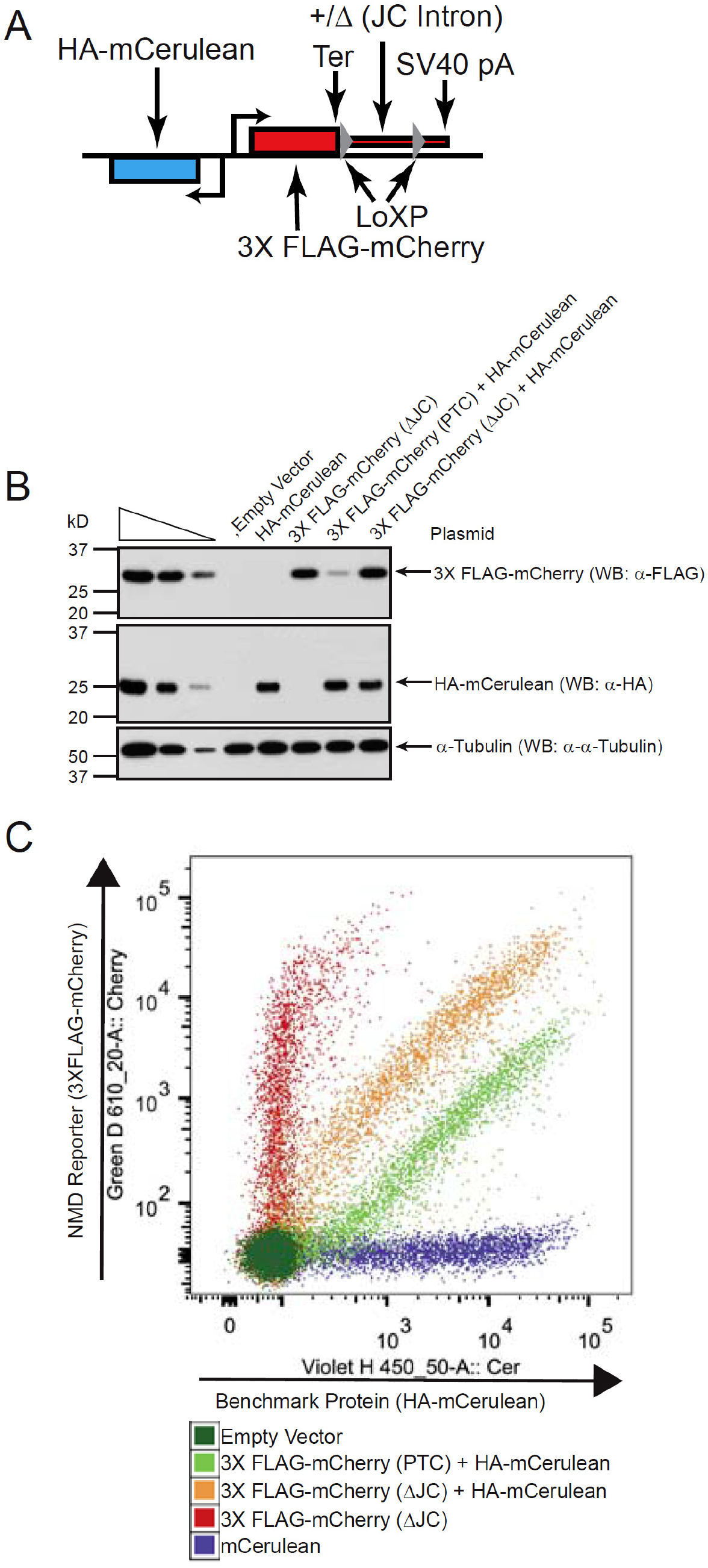
**Nonsense-mediated mRNA Decay (NMD) Reporter Design and Validation (A)** A bidirectional human cytomegalovirus immediate-early promoter drives production of HA-mCerulean in the sense direction (**r+**) and the NMD reporter in the antisense direction (**\*-**,). The NMD reporter is composed of the coding sequence for 3XFLAG epitope-tagged mCherry followed by a termination codon (Ter). Sequences 3′ to the termination codon derive from the TCRß gene and, in the +JC intron construct, contain the JC intron located >55-nucleotides downstream of the termination codon, forcing the termination codon to be recognized as a premature termination codon (PTC). TCRß-derived sequences are flanked by LoxP sites, allowing the JC intron to be deleted by application of Cre-Adenovirus. An SV40 polyadenylation site (pA) lies downstream of the final LoxP site. As a control, an identical construct except for deletion of the JC intron (AJC) was generated, as were single-color plasmids bearing only the HA-mCerulean or the 3XFLAG-mCherry (AJC) genes. **(B)** HEK293T cells were transiently transfected with the indicated plasmid and, 24-hours later, cells were subjected to western blotting (WB) using the specified antibodies (a) to assay for the presence of the indicated proteins. The three leftmost lanes under the wedge consist of 3-fold dilutions of lysate demonstrating that analyses are in the linear range of analysis. The level of cellular a-Tubulin serve as a loading control. kD, kilodaltons. **(C)** HEK293T cells from (B) were subjected to flow cytometry, as indicated. Green D610_20-A::Cherry refers to signal from a Green 532 nm laser with a 610/20 nm band pass filter collecting signal from the mCherry fluorescence. Similarly, Violet H450_50-A::Cer refers to signal from a Violet407 nm laser with a 450/50nm band pass filter collecting signal from the mCerulean fluorescence.

Transcription of the mCherry NMD reporter is driven by a bidirectional reporter derived from the pBi-CMV1 vector (Clontech) with the NMD reporter in the antisense direction (see below). In the sense direction, we cloned the fluorescent protein mCerulean, tagged at its N-terminus with an HA epitope, as an internal control. To deliver these reporters to HAP1 cells as vesicular-stomatitis virus glycoprotein (VSV-G)-pseudotyped viral particles, these constructs were cloned into an HIV-1-based lentiviral vector so as to avoid mapping reporter integration sites after the mutagenesis step, which is mediated by integration of a murine stem-cell virus (MSCV)-based retroviral gene-trap virus. Cloning the intron-containing reporter in the antisense direction relative to the lentiviral long terminal repeats (LTRs) limits truncation of the viral genome (and the consequential low viral titers) during virus production. As compensation controls for FACS, we also generated single-color versions of our reporters, encoding either the ΔJC mCherry reporter or the mCerulean protein. Transient transfection of HEK293T cells with these constructs showed that the mCherry NMD reporter protein was adequately expressed for biochemical analysis (**Fig. 1B**) as well as FACS analysis (**Fig. 1C**). Importantly, deletion of the cis-information that directs the reporter transcript to NMD, i.e. deletion of the JC intron, yielded an ~10-fold increase in both protein and fluorescence in this transient transfection assay.

### Reporter Cell Line Screening and Validation

HAP1 cells were infected with lentiviral particles encoding our reporter constructs and, in parallel, the control constructs. Considering that the position of integration can affect construct expression, we screened monoclonal populations of infected HAP1 cells for a reporter cell line that showed an adequate signal-to-noise ratio. The cell lines we sought should satisfy two criteria: first, they should show adequate mCherry signal to be detectable by FACS without further perturbation and second, the mCherry fluorescence should increase when NMD is disrupted. Since HAP1 cells are difficult to transfect, to mimic the situation in the second criteria, we delivered Cre-Adenovirus to monoclonal cell populations bearing the NMD reporter. This both deletes the TCRß-based 3′UTR region of the reporter, including the JC intron, rendering the transcript produced immune to NMD and also allows individual monoclonal populations to be screened for cell lines showing an adequate increase in fluorescence. We conducted two successive rounds of screening, arriving at our final reporter cell line (P6_9_E12). Adeno-Cre-mediated deletion of the JC intron increased the amount of protein produced from the mCherry reporter (**Fig. 2A**) as well as increased the fluorescence by FACS (**Fig. 2B**).

**Figure 2.**
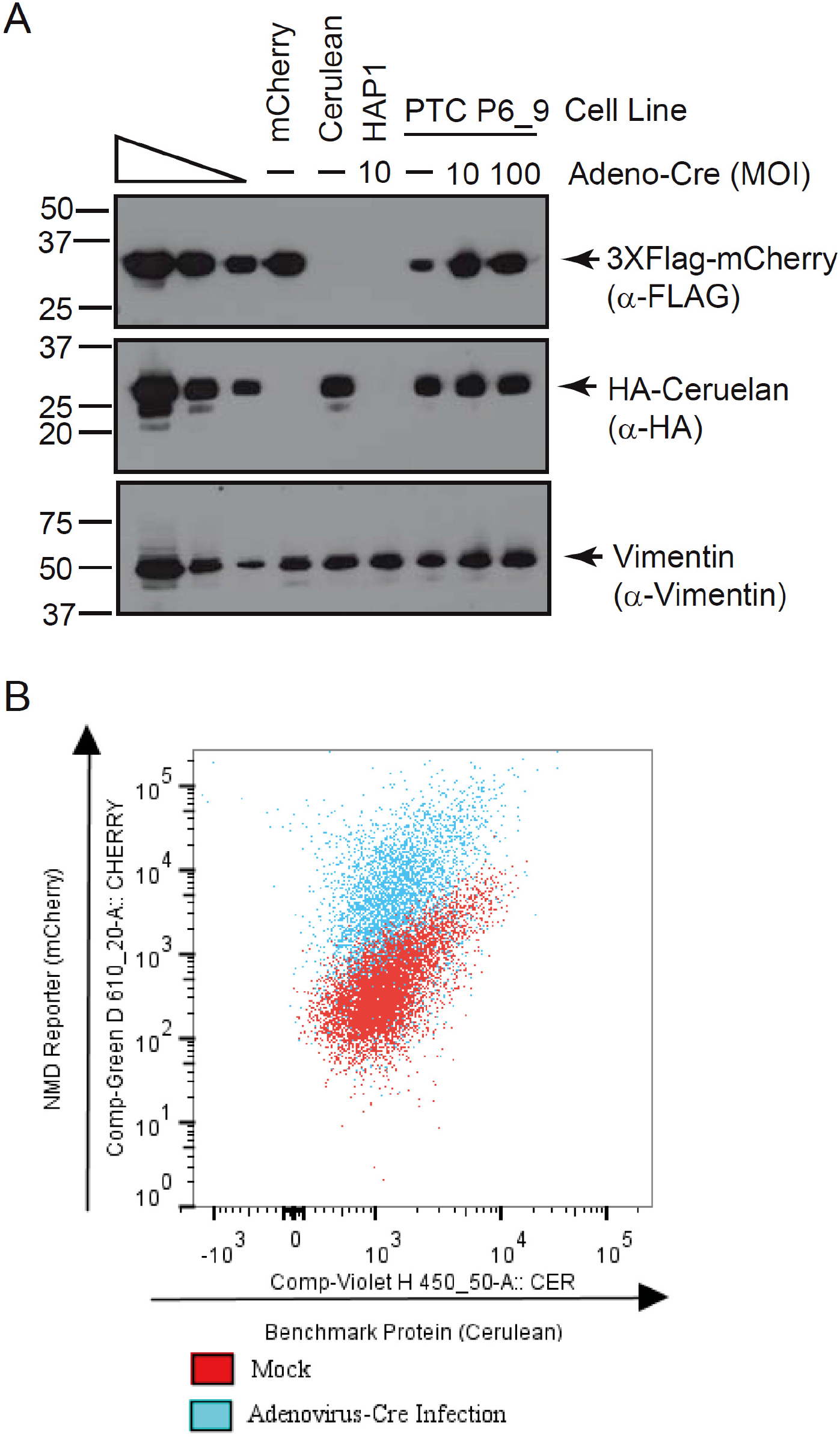
**HAP1 Stable Cell Line Validation (A)** HAP1 cell lines stably expressing the indicated protein (the PTC P6_9 cell line expresses both HA-mCerulean as well as 3XFLAG-mCherry(PTC)) were infected with Cre-Adenovirus (Adneo-Cre) at the indicated multiplicity of infection (MOI) to delete TCRß-derived sequences including the JC intron, so as to stabilize (i.e. obviate the NMD of) 3XFLAG-mCherry(PTC) mRNA. Cells were subjected to western blotting using the indicated antibodies (a). **(B)** PTC6_9 HAP1 cells from (A) were either mock infected or infected with Cre-Adenovirus as indicated in (A). Cells were then subjected to flow cytometry. Green D610_20-A::Cherry refers to signal from a Green 532 nm laser with a 610/20 nm band pass filter collecting signal from the mCherry fluorescence. Similarly, Violet H450_50-A::Cer refers to signal from a Violet407 nm laser with a 450/50nm band pass filter collecting signal from the mCerulean fluorescence.

### Retroviral Gene-Trap Mutagenesis

As a mutagen, we used an extensively described (23) gene-trap retrovirus. This virus encodes a strong portable splice acceptor site taken from the adenovirus serotype 40 long fiber protein (33), followed by the enhanced green fluorescent protein (eGFP) coding region and the SV40 polyadenylation site. Should the gene trap virus integrate into a host gene with its splice acceptor site in the sense orientation relative to the rest of the gene, an aberrant mRNA will be produced. Any host gene exons residing 5′ of the gene-trap integration site are spliced to the splice acceptor site, producing a non-functional eGFP fusion protein that is likely unstable. The gene-trapping method has been used extensively to generate null alleles in mice (34,35). We produced VSV-G pseudotyped gene-trap retroviral particles and infected ~100 million HAP1 cells. Greater than 90% of the cells obtained a gene-trap integration, as assessed by flow cytometry (**Fig. 3A**).

**Figure 3.**
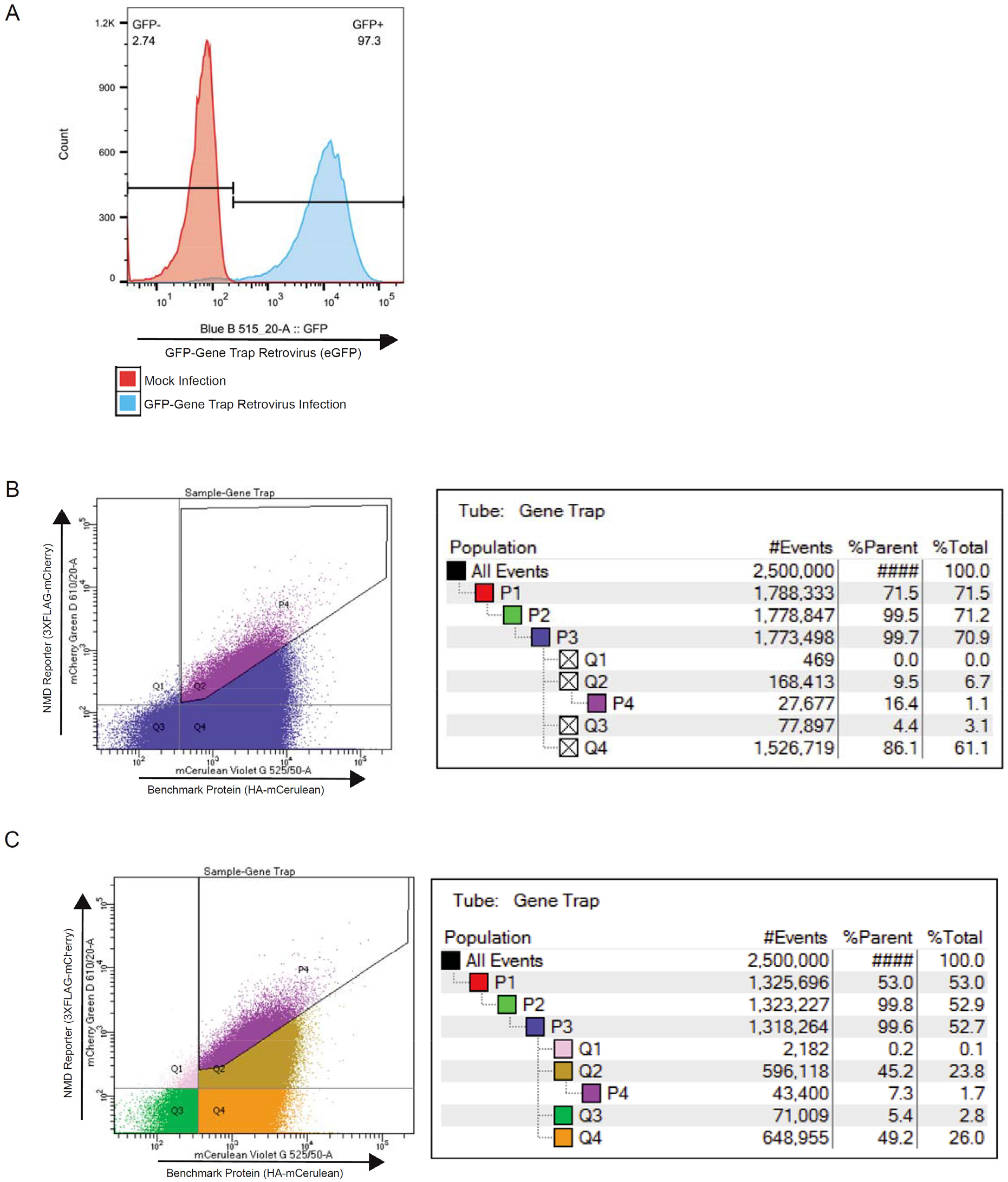
**eGFP Gene-Trap Retrovirus Infection (A)** P6_9_E12 HAP1 cells expressing the NMD reporter were either mock-infected or infected with the eGFP gene-trap retrovirus. Cells were then subjected to flow cytometry to estimate the percentage of eGFP-positive (i.e. infected) cells. Blue B 515 20-A::GFP refers to signal from a blue 488 nm laser with a 515/20 nm band pass filter collecting signal from eGFP fluorescence. **(B)** Infected P6_9_E12 cells from (A) were subjected to cell sorting, and the top 1-3% of mCherry-expressing cells were collected. (B) is representative of multiple sorts that were performed to amass a total of ~10 million collected cells. Statistics and cell counts for each quadrant (Q1, Q2, Q3, Q4) and the sort gate (P4) displayed on the FACS plot (left) are given in table format (right). Gates P1, P2, and P3 (not shown) were used to gate for whole cells (P1), and single events (P2 and P3) using forward and side-scatter. **(C)** A second iterative sort was performed. Cells from (B) were pooled, expanded, and subjected to cell sorting to again collect the top 1-3% of mCherry-expressing cells. (C) is representative of multiple sorts that were performed to amass a total of ~10 million collected cells, as described in (B). Statistics and cell counts for each quadrant (Q1, Q2, Q3, Q4) and the sort gate (P4) displayed on the FACS plot (left) are given in table format (right). Gates P1, P2, and P3 (not shown) were used to gate for whole cells (P1), and single events (P2 and P3) using forward and side-scatter.

### Selection and Sequencing

Gene-trap virus integration into genes encoding NMD trans-effectors will dampen NMD activity, leading to an increase in mCherry production and fluorescence, as mimicked by deletion of the JC intron in our controls (**Fig. 2B**). Thus we isolated by FACS ~10 million cells representing ~1-3% of cells with the highest mCherry expression in our mutagenized population (**Fig. 3B**). These cells were expanded, and a second round of sorting, taking the top ~1-3% of mCherry-expressing cells, was performed to further enrich for cells harboring insertions in potential NMD trans-effector genes (**Fig. 3C**).

With both our selected population enriched for mutations in genes encoding proteins important for NMD and our unselected mutagenized population in hand, we identified gene-trap insertion sites by high-throughput sequencing. Using a biotinylated primer annealing to the gene-trap retrovirus, we employed a linear amplification-mediated (LAM) PCR protocol (22) to amplify fragments corresponding to the genomic locus in which the gene-trap retrovirus integrated. After purification of these fragments using streptavidin-decorated magnetic dynabeads, a phosphorylated linker oligonucleotide was ligated to their 3′ ends using the commercially available CircligaseII enzyme. This was followed by a second exponential PCR step to add the Illumina P5 and P7 sequences. Products from the final PCR reactions should generate a homogenous smear when visualized in an agarose gel—discrete bands indicate that one PCR product has dominated the reaction, and these reactions should be discarded (**Fig. 4A**). Homogenous PCR reactions were pooled and sequenced using an Illumina HiSeq2500 instrument in single read mode and a custom sequencing primer that anneals to the very 5′ end of the gene-trap retrovirus 5′ LTR.

**Figure 4.**
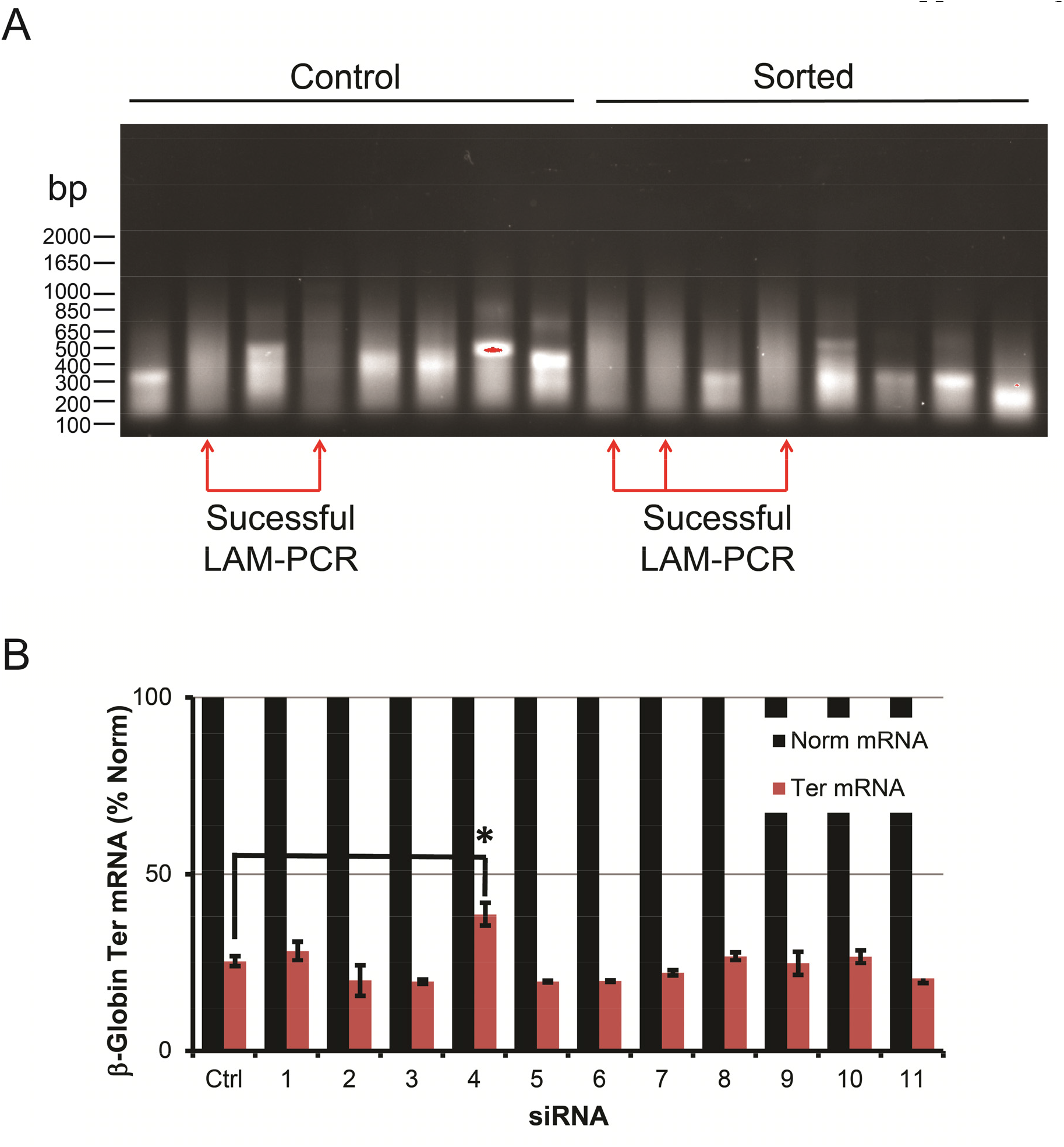
**LAM-PCR and Hit Validation (A)** LAM-PCR was performed on sorted cells as well as unsorted, control cells. Reactions were analyzed in 1% agarose gels, and only reactions showing homogenous smears (indicated by red arrows) were pooled and subjected to parallel deep-sequencing. bp, base-pairs. **(B)** Potential hits from the haploid screens were verified using an independent method that assayed for NMD in HEK 293T cells. Cells were transfected with one of 11 siRNAs representing 9 potential hits and, 24-hours later, transfected with either a ß-globin Norm test plasmid and mouse urinary protein (MUP) reference plasmid, or a ß-globin Ter test plasmid bearing a PTC at nucleotide 39 and the MUP reference plasmid. RNA was extracted, and reverse transcription-linked to quantitative PCR (RT-qPCR) was used to quantitate the levels of ß-globin and MUP mRNAs. The level of each ß-globin mRNA was normalized to the level of MUP mRNA, to control for variations in transfection efficiencies and RNA recovery. For each siRNA, the normalized level of ß-globin Ter mRNA was then defined as a percentage of the level of the normalized level of ß-globin Norm mRNA, the latter of which was defined as 100. Results were performed in biological triplicate. Norm mRNA, ß-globin Norm mRNA, which is not subjected to NMD; Ter mRNA, ß-globin Ter mRNA, which is subjected to NMD; Ctrl; control siRNA; *, p=0.010.

### Data Processing

Data were processed using a pipeline available at: https://github.com/BrummelkampResearch. Since each insertion event is sequenced multiple times, reads must be consolidated and then aligned to the human genome (hg19). Insertion sites were then mapped to intragenic regions, discarding insertions that were antisense to a gene, within a 3′ UTR, or within regions of overlapping genes. Finally, the number of unique disruptive insertions within each gene was tabulated for each population of cells (either unselected control cells, or FACS-sorted cells) as were the total number of disruptive insertions within each population. The ratio of these two numbers gives a metric that can be compared between populations for each gene using a one-sided Fisher’s exact test, and P-values are false discovery rate (FDR)-corrected (Benjamini-Hochberg). Potential hits were then ranked by P-value.

### Hit Validation

We verified hits using a previously described ß-globin mRNA-based reporter and human embryonic kidney (HEK)293T cells. HEK293T cells were first transfected with either a control non-targeting siRNA or one of a number of siRNAs corresponding to high P-value targets from our haploid genetic screen. Cells were then transfected 24-hours later with either a ß-globin Norm test plasmid and a mouse urinary protein (MUP) reference plasmid, or separately, a ß-globin Ter test plasmid bearing a PTC at nucleotide 39 and the MUP plasmid. mRNA levels were assessed by reverse transcription coupled to quantitative PCR (RT-qPCR). For each transfection, ß-globin mRNA levels were normalized to MUP mRNA levels to control for variations in transfection efficiencies and in RNA recovery. For each individual siRNA, the level of normalized ß-globin Ter mRNA was expressed as a percentage of the level of normalized ß-globin Norm mRNA. Analysis of 11 siRNAs against 9 proteins revealed that the depletion of one protein significantly increased the amount of ß-globin Ter mRNA relative to control siRNA (**Fig. 4B**), showing the utility of the haploid genetic screening method for uncovering novel proteins that modulate NMD activity.

## Discussion

Here, we have outlined how a haploid genetic screen selecting for gene mutations that decrease NMD activity was designed and implemented. Our goal is to enrich the existing literature of published haploid screens (22-24,26-30) by providing additional details in our protocol that should assist other investigators in implementing screens based on the selection intracellular fluorescence-based reporters. The full results of this screen and investigation into the mechanistic details of how confirmed hits modulate NMD activity will be reported elsewhere.

## Materials and Methods

### Cells and Cell Culture

Near-haploid HAP1 cells (25) were cultivated in Iscove‘s Modified Dulbecco’s Medium (Gibco) with 10% fetal bovine serum (FBS). HEK293T (ATCC) and Lenti-X HEK 293T cells (Clontech) were cultivated in Dulbecco’s Modified Eagle Medium (Gibco) with 10% FBS. HEK 293T cells for gene trap retrovirus production were cultivated in DMEM with 30% FBS.

### Plasmid Constructs

3XFLAG-mCherry(PTC) with HA-mCerulean, 3XFLAG-mCherry(ΔJC), 3XFLAG-mCherry(ΔJC), and HA-mCerulean plasmids were constructed as previously described (7). ß-globin and MUP plasmids were also previously described (7).

### Virus Production, Cell Infection, and FACS Sorting

For production of lentivirus, Lenti-X HEK293T cells were used with Lenti-X VSV-G packaging single shots (Clontech) according to manufacturer’s directions. Infected HAP1 cells were selected with Zeocin (200 μg/ml), followed by single-cell sorting into a 96-well plate using a BD LSR-II machine. Colonies were expanded, split into two plates, and then one plate was infected with Ad5CMV-Cre virus (University of Iowa Gene Transfer Vector Core) at an MOI of 10. mCherry and mCerulean fluoresence of infected and mock-infected cells was recorded by flow cytometry (BD, LSR-II machine), and colonies displaying a large increase in fluorescence after infection were further expanded and subjected to another round of screening by infection with Ad5CMV-Cre.

For production of eGFP gene-trap retrovirus (23), producer HEK 293T cells were cultured in DMEM supplemented with 30% FBS and plated into T-175 flasks 16-hours before transfection such that they were 80% confluent at transfection. Twelve flasks were used per virus preparation. For each flask, cells were transfected with 6.02 μg pCG-GAG/Pol, 1.4 μg pCMV-VSV-G, 0.84 μg pAdvantage (Promega), 2.2 μg pGT GFP(0), 2.2 μg pGT GFP (+1), and 2.2 μg pGT GFP (+2) using 45 μl Turbofectin 8.0 (Origene) in 850 μl Optimem (Gibco). Viral supernatants were harvested at 40 hours and then every 24-hours thereafter for a total of four harvests. Virus was pelleted by ultracentrifugation in an SW-28 rotor (Beckman) at 21,000 rpm for 2 hours at 4oC. Viral pellets were resuspended in a minimal amount of Hanks Buffered Saline Solution (HBSS, Gibco). Sixty million P9_9_E12 HAP1 cells were plated at 7.5 million cells per 15-cm dish 24-hours before infection with the gene-trap retrovirus. For infection, concentrated viral supernatant was applied to cells with 10μg/ml protamine sulfate (MP Biomedicals). Infection was performed 4 times at 24-hour intervals. Infected cells were then frozen at 10 million cells per vial for temporary storage at -80oC.

Infected cells were then subjected to FACS sorting using a BD LSR-II machine to isolate the top 1-3% of mCherry-expressing cells. Ten million cells were isolated in this way, expanded, and subjected to a second round of FACS sorting.

### LAM-PCR and Sequencing

LAM-PCR was performed essentially as described (22), with the following additional details. Genomic DNA from 10 million sorted and 10 million unsorted control cells was isolated using a QIAamp DNA mini kit (Qiagen) according to manufacturer’s directions, except elution was performed using 110 μl of H_2_O after a 15-minute incubation at 37oC. This was done twice, followed by elution using 60 μl of H_2_O and a 15-minute incubation for a total of 270 μl of genomic DNA at a concentration of ~200ng/μl. Sixteen LAM-PCR reactions for each population (sorted or control unsorted) were performed, with 2 μg of input DNA each and 0.375 pmol of double-biotinylated primer per reaction.

PCR reactions were pooled (2 LAM-PCR reactions per tube), and biotinylated DNA was purified using 20 μl of M-270 Streptavidin-coupled Dynabeads (Life Technologies), non-stick RNase-free microfuge tubes (Ambion), and 1:1 volume of 2X lithium binding buffer (6M LiCl, 10 mM Tris, 1 mM EDTA, pH 7.5). Binding was performed overnight at 4oC, followed by 4 washes with 800 μl of PBS/0.05% Triton-X 100.

Linker ligation to the 3′ end of PCR products was performed using Circligase II (Lucigen) exactly as described (22). Beads were then washed 4 times with 800 μl of PBS/0.05% Triton-X 100.

For the second, exponential PCR reaction, beads containing 2 LAM-PCR reactions per tube (i.e., one linker ligation reaction) were used for one exponential PCR reaction. PCR was performed exactly as described (22). PCR reaction supernatants were analyzed in a 1% agarose gel, then separately purified using a QIAquick PCR Purification kit (Qiagen). Reactions showing homogenous smears only (**Fig. 4A**) were pooled and submitted for sequencing on a HiSeq2500 instrument (Illumina) as described (22). Sequencing was performed by the University of Rochester Functional Genomics Center. Data were analyzed using a standard pipeline available at https://github.com/BrummelkampResearch.

### Immunoblotting

Immunoblotting was performed as described (7).

### Hit Validation

HEK293T cells were plated at 41,570 cells per well of a 24-well plate 24-hours before transfection. Cells were then transfected with 12.31 pmol of siRNA (Dharmacon or Sigma) per well using 0.6155 μl RNAiMAX (Invitrogen). Cells were transfected 24-hours later with either 17.687 ng of pCMV-ß-globin Norm and 7.075 ng of pCMV-MUP or, separately, with 17.687 ng of pCMV-ß-globin 39 Ter and 7.075 ng of pCMV-MUP plasmid. Cells were harvested 24-hours later. RNA was extracted and subjected to reverse transcription and qPCR as described (7).

## Acknowledgements

We thank Joppe Nieuwenhuis and Thijn Brummelkamp for reagents and helpful advice.

## References

1 Popp, M. W., and Maquat, L. E. (2013). Annu Rev Genet 47, 139-139.

2 Gardner, L. B. (2008). Mol Cell Biol 28, 3729-3729.

3 Gardner, L. B. (2010). Mol Cancer Res 8, 295-295.

4 Jia, J., Furlan, A., Gonzalez-Hilarion, S., Leroy, C, Gruenert, D. C, Tulasne, D., and Lejeune, F. (2015). Cell Death Differ 22, 1754-1763.

5 Karam, R, Wengrod, J., Gardner, L. B., and Wilkinson, M. F. (2013). Biochim Biophys Acta 1829, 624-624.

6 Nasif, S., Contu, L., and Mühlemann, O. (2018). Semin Cell Dev Biol 75, 78-78.

7 Popp, M. W., and Maquat, L. E. (2015). Nat Commun 6, 6632.

8 Popp, M. W., and Maquat, L. E. (2018). Curr Opin Genet Dev 48, 44-44.

9 Wang, D., Zavadil, J., Martin, L., Parisi, F, Friedman, E., Levy, D., Harding, H., Ron, D., and Gardner, L. B. (2011). Mol Cell Biol 31, 3670-3670.

10 Yamashita, A., Izumi, N., Kashima, I., Ohnishi, T., Saari, B., Katsuhata, Y., Muramatsu, R., Morita, T., Iwamatsu, A., Hachiya, T., Kurata, R., Hirano, H., Anderson, P., and Ohno, S. (2009). Genes Dev 23, 1091-1091.

11 Anastasaki, C., Longman, D., Capper, A., Patton, E. E., and Cáceres, J. F. (2011). Nucleic Acids Res 39, 3686-3686.

12 Longman, D., Hug, N., Keith, M., Anastasaki, C., Patton, E. E., Grimes, G., and Cáceres, J. F. (2013). Nucleic Acids Res 41, 8319-8319.

13 Alexandrov, A., Colognori, D., Shu, M. D., and Steitz, J. A. (2012). Proc Natl Acad Sci U S A 109, 21313-21313.

14 Baird, T. D., Cheng, K. C., Chen, Y. C., Buehler, E., Martin, S. E., Inglese, J., and Hogg, J. R. (2018). Elife 7.

15 Ge, Z., Quek, B. L., Beemon, K. L., and Hogg, J. R. (2016). Elife 5.

16 Nickless, A., Jackson, E., Marasa, J., Nugent, P., Mercer, R. W., Piwnica-Worms, D., and You, Z. (2014). Nat Med 20, 961-961.

17 Karam, R., Lou, C. H., Kroeger, H., Huang, L., Lin, J. H., and Wilkinson, M. F. (2015). EMBO Rep 16, 599-599.

18 Lou, C. H., Dumdie, J., Goetz, A., Shum, E. Y., Brafman, D., Liao, X., Mora-Castilla, S., Ramaiah, M., Cook-Andersen, H., Laurent, L., and Wilkinson, M. F. (2016). Stem Cell Reports 6, 844-844.

19 Lou, C. H., Shao, A., Shum, E. Y., Espinoza, J. L., Huang, L., Karam, R., and Wilkinson, M. F. (2014). Cell Rep 6, 748-748.

20 Alexandrov, A., Shu, M. D., and Steitz, J. A. (2017). Mol Cell 65, 191-191.

21 Blomen, V. A., Majek, P., Jae, L. T., Bigenzahn, J. W., Nieuwenhuis, J., Staring, J., Sacco, R., van, Diemen F. R., Olk, N., Stukalov, A., Marceau, C., Janssen, H., Carette, J. E., Bennett, K. L., Colinge, J., Superti-Furga, G., and Brummelkamp, T. R. (2015). Science 350, 1092-1092.

22 Brockmann, M., Blomen, V. A., Nieuwenhuis, J., Stickel, E., Raaben, M., Bleijerveld, O. B., Altelaar, A. F. M., Jae, L. T., and Brummelkamp, T. R. (2017). Nature 546, 307-307.

23 Carette, J. E., Guimaraes, C. P., Varadarajan, M., Park, A. S., Wuethrich, I., Godarova, A., Kotecki, M., Cochran, B. H., Spooner, E., Ploegh, H. L., and Brummelkamp, T. R. (2009). Science 326, 1231-1231.

24 Carette, J. E., Guimaraes, C. P., Wuethrich, I., Blomen, V. A., Varadarajan, M., Sun, C., Bell, G., Yuan, B., Muellner, M. K., Nijman, S. M., Ploegh, H. L., and Brummelkamp, T. R. (2011). Nat Biotechnol 29, 542-542.

25 Carette, J. E., Pruszak, J., Varadarajan, M., Blomen, V. A., Gokhale, S., Camargo, F. D., Wernig, M., Jaenisch, R., and Brummelkamp, T. R. (2010). Blood 115, 4039-4039.

26 Carette, J. E., Raaben, M., Wong, A. C., Herbert, A. S., Obernosterer, G., Mulherkar, N., Kuehne, A. I., Kranzusch, P. J., Griffin, A. M., Ruthel, G., Dal Cin, P., Dye, J. M., Whelan, S. P., Chandran, K., and Brummelkamp, T. R. (2011). Nature 477, 340-340.

27 Nieuwenhuis, J., Adamopoulos, A., Bleijerveld, O. B., Mazouzi, A., Stickel, E., Celie, P., Altelaar, M., Knipscheer, P., Perrakis, A., Blomen, V. A., and Brummelkamp, T. R. (2017). Science 358, 1453-1453.

28 Staring, J., van den Hengel, L. G., Raaben, M., Blomen, V. A., Carette, J. E., and Brummelkamp, T. R. (2018). Cell Host Microbe 23, 636-643 e635.

29 Staring, J., von Castelmur, E., Blomen, V. A., van den Hengel, L. G., Brockmann, M., Baggen, J., Thibaut, H. J., Nieuwenhuis, J., Janssen, H., van, Kuppeveld F. J., Perrakis, A., Carette, J. E., and Brummelkamp, T. R. (2017). Nature 541, 412-412.

30 Jae, L. T., Raaben, M., Riemersma, M., van Beusekom, E., Blomen, V. A., Velds, A., Kerkhoven, R. M., Carette, J. E., Topaloglu, H., Meinecke, P., Wessels, M. W., Lefeber, D. J., Whelan, S. P., van Bokhoven, H., and Brummelkamp, T. R. (2013). Science 340, 479-479.

31 Paillusson, A., Hirschi, N., Vallan, C., Azzalin, C. M., and Mühlemann, O. (2005). Nucleic Acids Res 33, e54.

32 Fiebiger, E., Hirsch, C., Vyas, J. M., Gordon, E., Ploegh, H. L., and Tortorella, D. (2004). Mol Biol Cell 15, 1635-1635.

33 Carette, J. E., Graat, H. C., Schagen, F. H., Abou El Hassan, M. A., Gerritsen, W. R., and van Beusechem, V. W. (2005). J Gene Med 7, 1053-1053.

34 Yamamura, K., and Araki, K. (2008). Cancer Sci 99, 1-1.

35 Araki, M., Araki, K., and Yamamura, K. (2009). Curr Pharm Biotechnol 10, 221-221.

